# Cross comparison of imaging strategies of mitochondria in *C. elegans* during aging

**DOI:** 10.1101/2024.12.24.630282

**Authors:** Juri Kim, Naibedya Dutta, Matthew Vega, Andrew Bong, Maxim Averbuhk, Rebecca Aviles Barahona, Athena Alcala, Jacob T. Holmes, Gilberto Garcia, Ryo Higuchi-Sanabria

## Abstract

Mitochondria are double membrane-bound organelles with pleiotropic roles in the cell, including energy production through aerobic respiration, calcium signaling, metabolism, proliferation, immune signaling, and apoptosis. Dysfunction of mitochondria is associated with numerous physiological consequences and drives various diseases, and is one of twelve biological hallmarks of aging, linked to aging pathology. There are many distinct changes that occur to the mitochondria during aging including changes in mitochondrial morphology, which can be used as a robust and simple readout of mitochondrial quality and function. Although mitochondrial morphology alone cannot be used to conclude the quality of mitochondria, it is highly correlated with mitochondrial function whereby mitochondria exhibit increased fragmentation with age in multiple cell types of the nematode *C. elegans.* Thus, *C. elegans* serve as a robust model for rapidly measuring mitochondrial morphology changes during aging. To standardize imaging methods for mitochondrial morphology in *C. elegans*, we provide a detailed comparative characterization of several transgenic constructs, highlighting benefits and caveats for aging biology studies.

**Summary Blurb:** This study evaluates mitochondrial imaging in *C. elegans* during aging, comparing various transgenic constructs for tissue-specific mitochondrial visualization. The findings highlight technical considerations, imaging method standardization, and the utility of *C. elegans* as a robust model for studying mitochondrial dynamics.

## Introduction

Mitochondrial fitness and function are critical for proper health and function of a cell due to their role in numerous cellular processes, including energy production, apoptotic and necrotic cell death regulation, calcium and amino acids storage, lipid oxidation, and heat production (Kamer & Mootha, 2015; Wang et al, 2023). Disrupting mitochondrial function can result in major consequences, including metabolic dysregulation, accumulation of toxic reactive oxygen species (ROS), and dysregulation of many cellular pathways (López-Otín et al, 2023; Zong et al, 2024). As such, mitochondrial dysfunction is one of twelve major biological hallmarks of aging, whereby mitochondrial dysfunction is observed during the natural aging process in most model organisms (López-Otín et al, 2023). Mitochondrial dysfunction can be defined by many measurable outcomes including loss of mitochondrial membrane potential, import, and respiratory capacity; accumulation of mitochondrial DNA mutations; loss of stoichiometry of multi-protein complexes in the mitochondria; and changes in mitochondrial morphology, mass, and volume (Amorim et al, 2022; Zong et al, 2024). Often, many of these features are correlated and occur simultaneously, which allows for the usage of one of these markers as a general readout for mitochondrial quality and function.

Mitochondrial morphology is a commonly used feature to indirectly determine mitochondrial function, as the quality control of mitochondria is regulated by mitochondrial dynamics, a tightly coordinated balance of continuous fusion and fission events that determine the shape, length, and number of mitochondria (Wang et al, 2023). The loss of balance between fusion and fission can have dramatic impacts on mitochondrial function, whereby excessive fission events or decline in fusion can lead to fragmentation of mitochondria, while excessive fusion or reduced fission can result in hyperfusion (de Boer et al, 2021). Generally, fragmentation and excessive fission of mitochondria is correlated with loss of membrane potential and loss of mitochondrial function (Zorova et al, 2018). This suggests that a shift towards fission is bad, and that more fusion would be beneficial; however, excessive mitochondrial fusion can disrupt important quality control machinery, such as mitophagy that clears damaged mitochondrial components (Ashrafi & Schwarz, 2013). Thus, it is the collective balance of fusion and fission events that are important in maintaining cellular homeostasis, and abnormal mitochondrial dynamics can result in pathology of age-related diseases including cardiovascular diseases (Wu et al, 2019; Quiles & Gustafsson, 2022), cancer (Ma et al, 2020), and lung disorders (Sharma et al, 2021). These studies highlight the importance of studying mitochondrial dynamics and visualizing their morphology during aging.

*Caenorhabditis elegans* serves as an excellent model system to study mitochondrial dynamics due to the low cost of their maintenance, established genome, and transparent body that allows for microscopic visualization of mitochondria in live animals. Importantly, their short lifespans allow for large-scale aging studies whereby mitochondrial imaging can be performed throughout the lifespan of the worm. Finally, there are robust genetic tools available for genetic modifications of *C. elegans* including CRISPR/Cas9 genome editing (Dickinson et al, 2015) and RNA interference (RNAi) (Bosher & Labouesse, 2000), which allow for identification of novel genetic mechanisms that impact mitochondrial dynamics and aging. Importantly, mitochondrial dynamics is highly conserved in *C. elegans* and the structure and function of mitochondria are highly similar to those of mammalian cells. Mitochondrial fusion in *C. elegans* is controlled by the conserved inner and outer membrane fusion proteins, EAT-3 (ortholog of Opa1) and FZO-1 (ortholog of Mfn1/2) (Rolland et al, 2009). Fission is controlled by the dynamin-related protein DRP-1 (ortholog of Drp1), which constricts the mitochondrion to separate mitochondrial membranes (Labrousse et al, 1999).

One of the most common methods to visualize mitochondria in *C. elegans* is to utilize genetically encoded mitochondria-localized fluorophores due to the ease of genetic manipulation in this model. However, many of the currently existing methods involve transgenic animals with high copy expression of fluorophores, including a mitochondrial matrix localized green fluorescent protein (hereafter referred to as MLS::GFP) (Hoffmann et al, 2009) or overexpression of a red fluorescent protein (RFP)-tagged mitochondrial-localized protein, such as TOMM-20 (Wang et al, 2018). The benefit of these high copy expression constructs is that since the fluorophores are expressed at very high levels and thus very bright, low sensitivity cameras and weak excitatory light sources could be used to robustly visualize mitochondria.

However, with the advent in technological advancement in microscopy and sensitivity of cameras in the past few decades, there is no longer a need for such high expression of fluorescent molecules for detection. Importantly, there are many potential caveats of high copy expression, including a potential stress to the mitochondria to import so many proteins into the mitochondria (Begelman et al, 2022). In fact, while preparing this manuscript, another group has independently identified that currently used methods suffer from several physiological caveats, including a significant reduction in lifespan (Valera-Alberni et al, 2024). In their manuscript, the Mair lab illustrates the advantages and disadvantages of currently available tools to image mitochondria and offer a suite of single-copy mitochondrial membrane-localized fluorophores and endogenously tagged mitochondrial proteins as alternative strategies.

In our study, we offer another alternative strategy for mitochondrial imaging using a single copy, matrix-localized fluorescent molecule. Here, we utilized mosSCI transgenics for precise, stable, and single-copy expression of MLS::GFP in a known genetic locus. We compare and contrast our imaging strategies with the most commonly used strains in an attempt to standardize methods to image mitochondria in the muscle, intestine, and hypodermis in *C. elegans*.

Importantly, our strains are complementary to the strains developed by the Mair lab and can be used to simultaneously visualize the outer membrane and mitochondrial matrix.

## Results

### Development of single-copy MLS::GFP strains using mosSCI in C. elegans

Mitochondrial morphology is often directly correlated with mitochondrial fitness and function and thus has gained popularity as the first line of study for understanding mitochondrial organization and quality under distinct circumstances. *C. elegans* serve as an exceptional model system for visualization of mitochondrial morphology, as the clear body allows for imaging of mitochondria in whole, live animals. However, current technologies for visualization of mitochondrial morphology have several distinct caveats: first, it utilized integration of a multi-copy MLS::GFP or TOMM-20::mRFP construct using a *myo-3* promoter for muscle-specific expression. These constructs are thus integrated in a random locus in the genome and have very high copy expression of these fluorescent proteins, which could potentially impact mitochondrial quality and organismal physiology. Indeed, a recent study showed that these strains had measurable changes in longevity, reproduction, animal size, and generation time (Valera-Alberni et al, 2024). Moreover, these animals exhibit highly variable expression in fluorescence across tissues even within the same animal, making comprehensive studies and quantitative imaging very challenging.

Here, we sought to make more simplified versions of these strains by expressing MLS::GFP using the mosSCI system (Frøkjær-Jensen et al, 2012) to eliminate several caveats of previously used methods. These animals have MLS::GFP integrated into a known locus, which simplifies genetic crosses and allows for controlled, equal expression of the fluorophore across the entire animal. By fusing the MLS of ATP-1 to GFP and using cell-type specific promoters, we created robust methods to visualize mitochondrial morphology specifically in the muscle (*myo-3p*), intestine (*vha-6p*), and the hypodermis (*col-19p*) (**Fig. 1A**).

**Fig 1.**
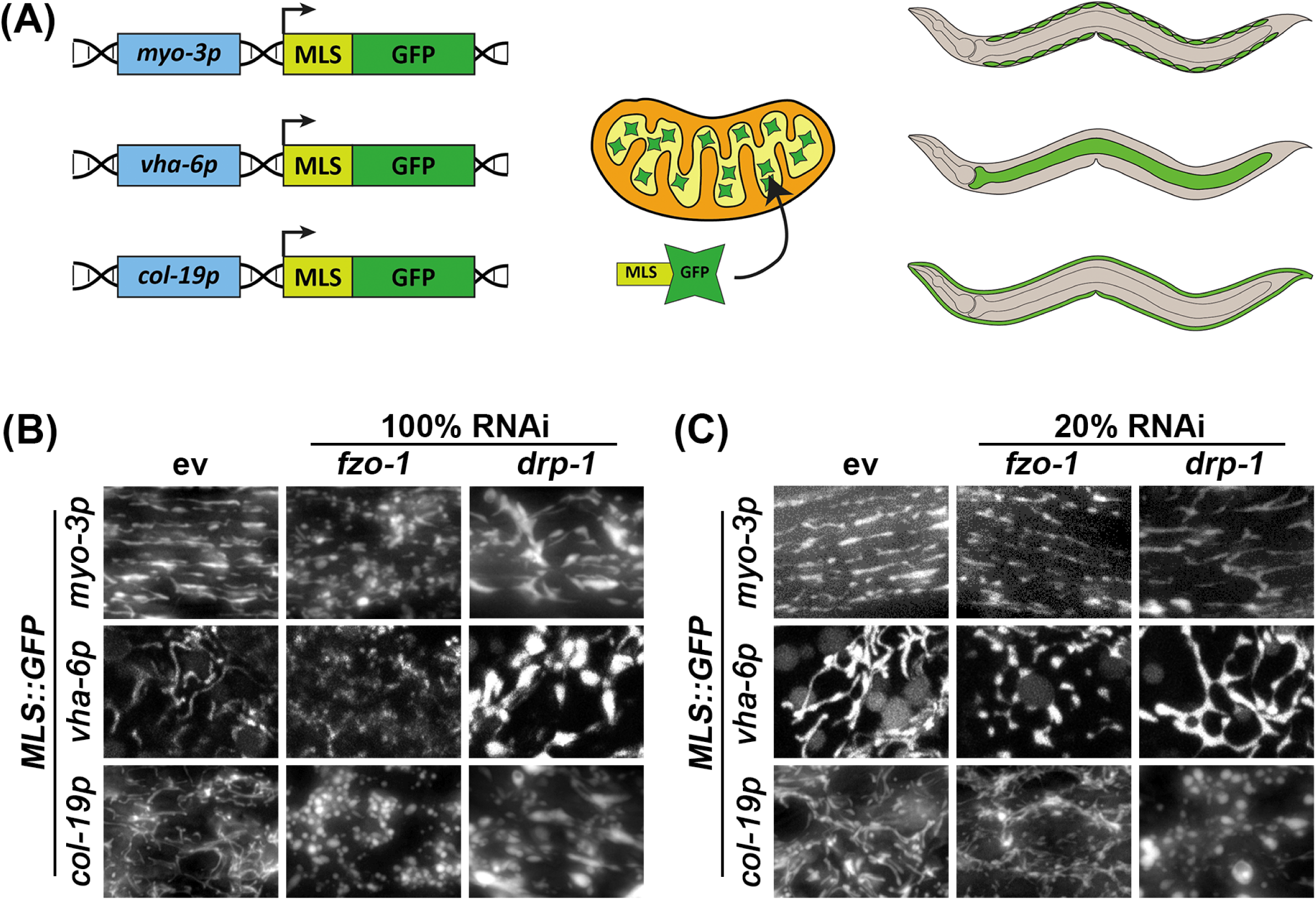
Validation of strains by altering mitochondrial morphology using *fzo-1* or *drp-1* RNAi treatments. (**A**) Schematic of MLS::GFP, which we express using cell-type specific promoters: *myo-3p* in the muscle, *vha-6p* in the intestine, and *col-19p* in the hypodermis. MLS::GFP is imported into the matrix of the mitochondria and can be robustly visualized in live animals using fluorescent microscopy. (**B**) Animals expressing cell-type specific MLS::GFP were grown on control empty vector (ev), *fzo-1*, or *drp-1* RNAi from the L1 stage and imaged on day 5 of adulthood. (**C**) *fzo-1* and *drp-1* RNAi were diluted to 20% with ev (i.e., 1:4 ratio of RNAi:ev). Animals were grown on the indicated RNAis from the L1 stage and imaged at day 5 of adulthood.

To test the dynamic range of these reporters for visualizing mitochondrial morphology, we exposed animals to RNAi knockdown of genes encoding the fusion and fission machinery, *fzo-1* and *drp-1*, respectively. As expected, knockdown of fusion resulted in significant fragmentation of mitochondria (**Fig. 1B**). However, RNAi knockdown of *drp-1* resulted in aggregation of mitochondria, which is consistent with previous findings (Labrousse et al, 1999) that argue that hyperfusion of mitochondria results in formation of aggregated mitochondria that resemble spheres (**Fig. 1B**). To better measure the dynamic range of fragmented versus fused mitochondria, we performed titration of *fzo-1* and *drp-1* knockdown and found that dilution of the *drp-1* RNAi with an empty vector (EV) RNAi to 20% (i.e., a 1:4 ratio of *drp-1:* EV) was optimal to block mitochondrial fusion and create a more tubular and interconnected structure in the muscle and intestine, rather than forming hyperfused spheres (**Fig. 1C**). However, in the hypodermis, 20% *drp-1* still resulted in mitochondrial spheres. A similar dilution of *fzo-*1 RNAi to 20% still effectively fragmented the mitochondria in all tissues, but to a lesser extent than undiluted RNAi (**Fig. 1C**). These data provide direct evidence that our mosSCI strains allow for robust visualization of mitochondrial morphology and behave as expected when mitochondrial dynamics are altered. Moreover, our data provide relative RNAi concentrations of *fzo-1* and *drp-1* that allow for alterations of mitochondrial morphology.

### Mitochondria exhibit fragmentation during aging

Next, we performed mitochondrial imaging during aging. Consistent with previous reports (Sharma et al, 2019), we see that animals display an increase in mitochondrial fragmentation during the aging process (**Fig. 2A**). Comparison of our mosSCI MLS::GFP strain to the most commonly used multi-copy integration muscle mitochondrial strains utilizing MLS::GFP and TOMM-20::mRFP showed that our strains exhibit a delayed fragmentation of mitochondria during aging (**Fig. 2B**). This is likely due to the potential detrimental effects of having to import a large quantity of mitochondria-localized proteins (Begelman et al, 2022). Similar to muscle mitochondrial imaging, high-copy expression of MLS::GFP in the intestine also resulted in premature mitochondrial fragmentation during aging compared to our mosSCI MLS::GFP strain (**Fig. 2C**).

**Fig. 2.**
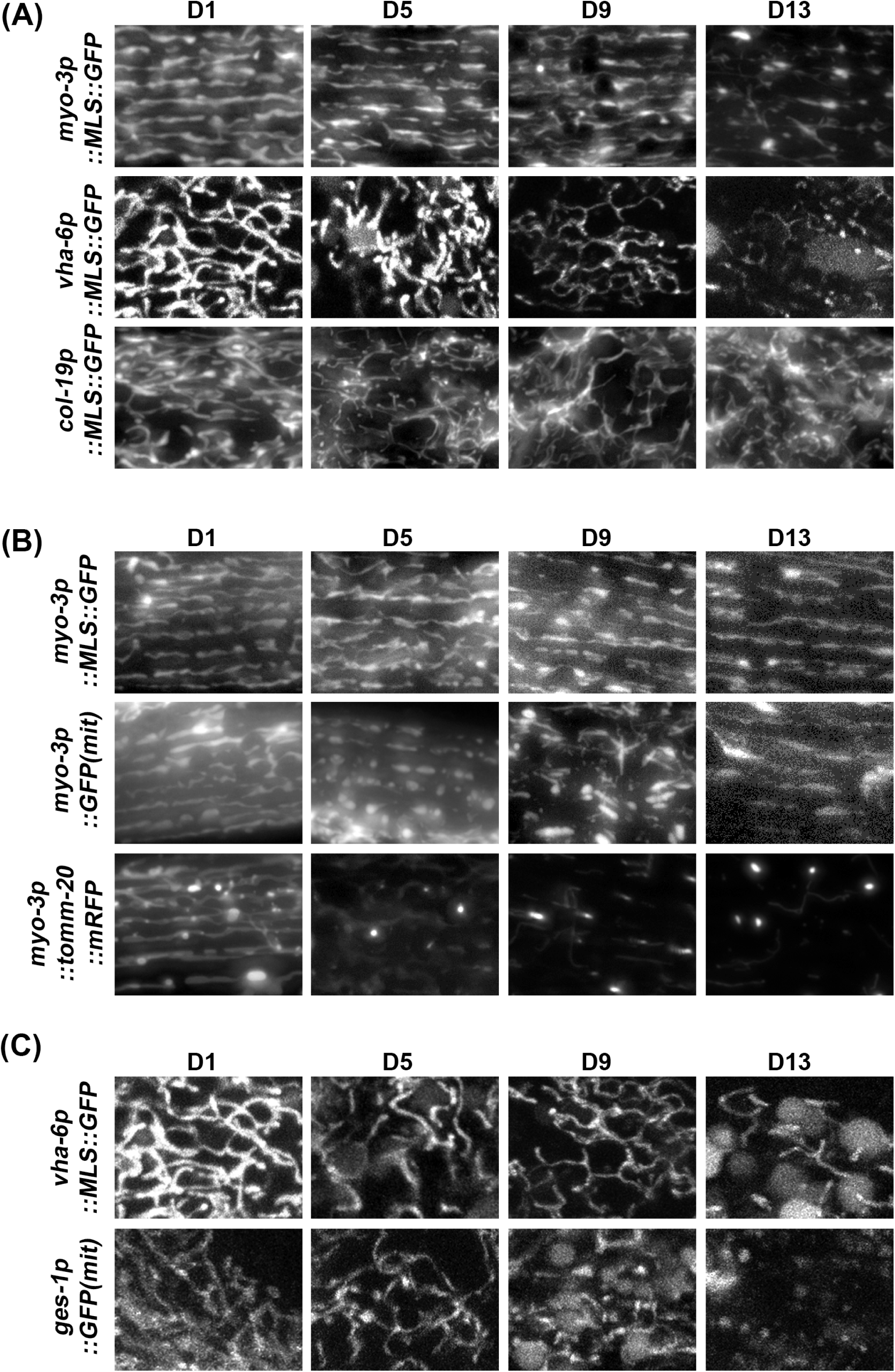
Comparison of strains for cell-type specific imaging of mitochondria during aging. **(A)** Imaging of cell-type specific mitochondria in the muscle (*myo-3p*), intestine (*vha-6p*), and hypodermis (*col-19p*) using a single-copy integration of MLS::GFP using mosSCI during aging. **(B)** Comparison of muscle mosSCI MLS::GFP (RHS191) strain to multi-copy *myo-3::GFP(mit)* (SJ4103) and multi-copy *myo-3p::TOMM-20::mRFP* (PS6192) during aging. (**C**) Comparison of intestine mosSCI MLS::GFP (RHS193) to multi-copy *ges-1p::GFP(mit)* (SJ4143) during aging. All animals were grown on ev from the L1 stage and imaged at day 1, 5, 9, and 13 of adulthood.

During the preparation of this manuscript, another group independently identified similar caveats of these previously validated strains used to visualize mitochondrial morphology (Valera-Alberni et al, 2024). Importantly, we confirmed the Mair lab’s findings that previously used strains exhibit significant variability in expression across cells, while our mosSCI strains showed consistent expression across cells (**Fig. S1A**). In this study, the Mair lab also created novel single-copy strains that utilize either a minimal MLS from the TOMM-20 protein fused to a fluorescent molecule or directly integrated a fluorescent tag at the endogenous gene locus of TOMM-50 or TIMM-70. Similar to the strains presented here, this study eliminated many of the caveats from previously utilized strains and are complementary to our imaging strategies. Since our fluorescent probes are localized to the mitochondrial matrix, it allows for imaging of the mitochondrial lumen, which can be directly paired with the outer membrane marker from the Mair lab to visualize multiple subcompartments of the mitochondria simultaneously (**Fig. S1B**). This is an important consideration for those interested in dynamics of inner and outer mitochondrial membrane fusion and fission, which does not always occur simultaneously (Malka et al, 2005). Indeed, we saw some instances where outer membrane RFP signal were detected in the absence of matrix GFP signal. An interesting phenomenon observed by the Mair lab was that when TOMM-20 or TIMM-50 were fused to RFPs, these proteins aggregated at old age, which did not occur when GFP was used. Similar to their findings, we also observe aggregation of the matrix-localized RFP, MLS::mRuby at old age, which we did not observe in any of our matrix-localized GFP strains (**Fig. S1C**).

While *C. elegans* offer a simple and easy way to study mitochondrial morphology during aging due to their short lifespan and ease of growth, one challenge is that they exist as hermaphrodites with the ability to self-fertilize. Therefore, for aging studies, progeny must be eliminated to prevent contamination of the aging cohort with their offspring. One common method of eliminating progeny is to chemically sterilize animals using exposure to 5-Fluoro-2′-deoxyuridine (FUDR), which causes developmental arrest in progeny by preventing DNA replication (Bell & Wolff, 1964; Rooney et al, 2014). However, FUDR may have unwanted effects on aging and some studies have shown that exposure to FUDR can potentially impact the aging process (Mitchell et al, 1979; Rooney et al, 2014). Therefore, we compared exposure to FUDR to the standard method of manually picking adults away from their progeny daily. Although we find that exposure to FUDR and manual picking of adults away from their progeny both show similar age-induced fragmentation of mitochondria, FUDR-exposed animals exhibited a slight delay in mitochondrial fragmentation in all cell types (**Fig. 3A-B**). Importantly, this difference was not entirely due to the manual manipulation of worms. Animals exposed to FUDR, but manually moved daily still exhibit a delay in mitochondrial fragmentation in muscle, but not in the intestine and hypodermis (**Fig. S2**). These data show that while exposure to FUDR may cause a minor delay in mitochondrial fragmentation with age in some tissue, there are no major artifacts induced by this aging method overall, and thus FUDR-exposure is likely a feasible approach to aging out animals for studies of mitochondrial morphology.

**Fig. 3.**
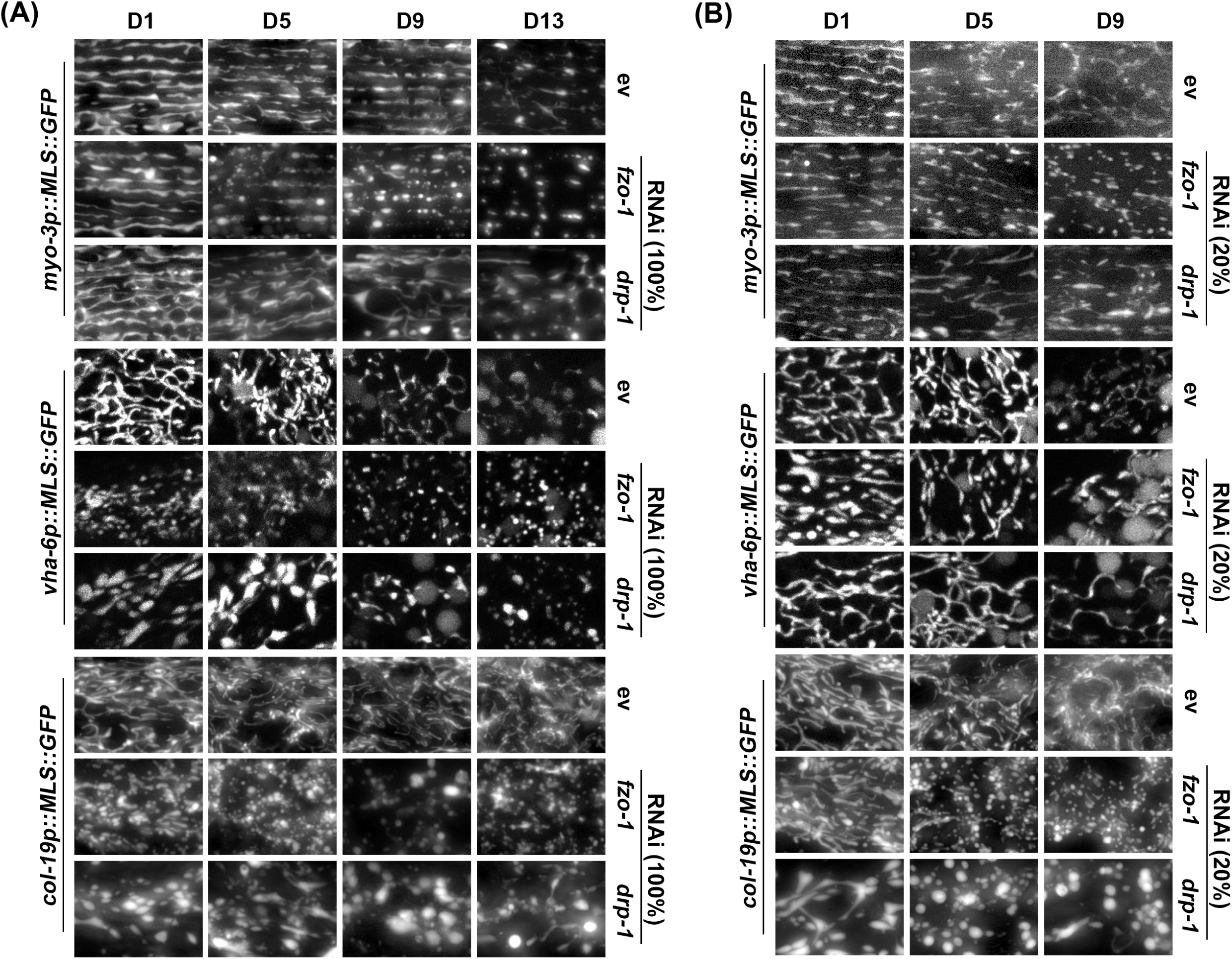
Imaging of mitochondrial morphology during aging upon *fzo-1* and *drp-1* RNAi. (**A**) Animals expressing MLS::GFP were grown on full concentration RNAi of control (ev), *fzo-1*, or *drp-1* from the L1 stage and imaged during day 1, 5, 9, and 13 of adulthood. (**B**) *fzo-1* and *drp-1* RNAi were diluted to 20% with ev (i.e., 1:4 ratio of RNAi:ev). Animals were grown on the indicated RNAis from the L1 stage and imaged at day 1, 5, 9, and 13 of adulthood.

As an alternative approach to aging animals for those who wish to avoid FUDR-exposure, another commonly used aging method includes using temperature-sensitive mutants including the germline mutant *glp-4(bn2)* (Beanan & Strome, 1992; Castro Torres et al, 2022) or the sperm-deficient mutants *fer-1* (Ward & Miwa, 1978) and *CF512* (Garigan et al, 2002). Here, the germline mutant *glp-4(bn2)* previously validated to not impact aging (TeKippe & Aballay, 2010) was crossed into our mosSCI MLS::GFP animals. Animals were grown at 22 °C for the duration of their lifespan, as this elevated temperature was sufficient to sterilize animals in our hands. We found that *glp-4(bn2)* displayed more pronounced fragmentation in all tissues at all timepoints even from early ages (**Fig. 3C**). Thus, care must be taken when using *glp-4(bn2)* animals for mitochondrial imaging studies.

Finally, we tested the impact of bacterial food source on mitochondrial morphology during aging. *E. coli* B strain OP50 and K strain HT115 are the most common food sources for *C. elegans* with OP50 being the most common food choice for standard maintenance and HT115 used for RNAi experiments (Reinke et al, 2010; Revtovich et al, 2019). However, previous work has shown that mitochondrial health is improved in worms fed an HT115 diet, likely due to an increased availability of vitamin B12 (Revtovich et al, 2019). This is an important consideration, since mitochondrial morphology can exhibit significant differences based on the bacterial diet (Revtovich et al, 2019; Neve et al, 2020). Consistent with previous reports, we found that animals grown on OP50 exhibit more fragmented mitochondrial morphology compared to animals grown on HT115 in the muscle, intestine, and hypodermis (**Fig. 4A**). Interestingly, many of these differences are most prominently observed at day 1 of adulthood, but differences were less noticeable during mid age and old age. To determine whether these differences were due to differences in vitamin B12 as previously described, we performed mitochondrial imaging in animals grown on OP50 diets supplemented with a vitamin B12 analog (adoCbl; adenosyl cobalamin, final concentration of 12.8 nM). We found that supplementation of adoCbl rescues the mitochondrial fragmentation in OP50 on day 1 of adulthood compared to HT115 food source in all tissues. Moreover, the supplementation was even shown to delay the age-associated mitochondrial fragmentation of old worms (day 9 of adulthood) both on HT115 and OP50. (**Fig. 4A-B**). Altogether, our data presented here provide evidence that our mosSCI generated strains are robust and reliable reporters for mitochondrial morphology during aging. More importantly, while some differences exist in terms of methodology for animal growth or aging, there are not dramatic differences between strategies and consistency in using a single method – or testing multiple methods – are both viable options for aging experiments.

**Fig. 4.**
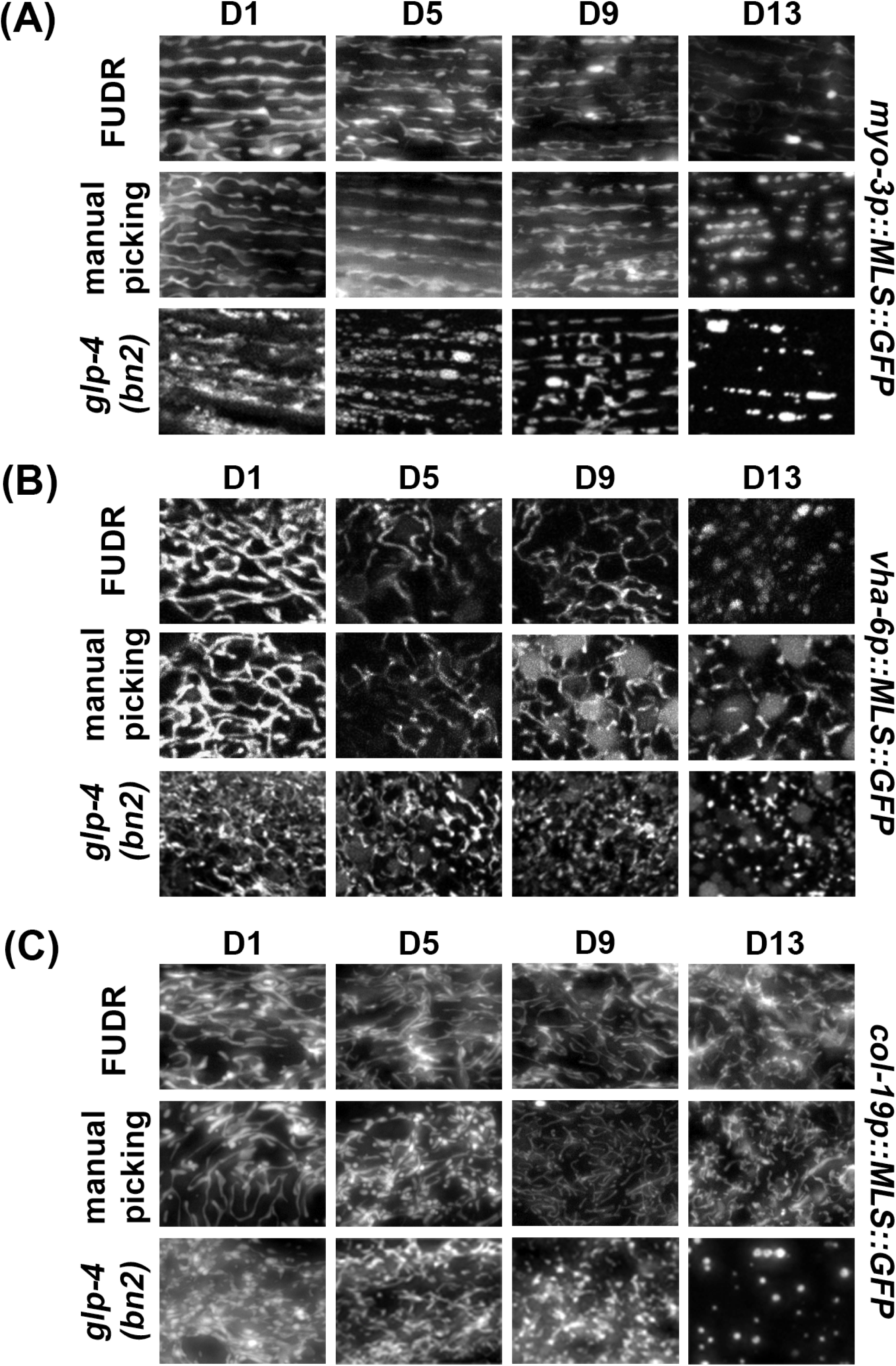
Comparison of aging methods for imaging of mitochondrial morphology. Animals were aged using the following methods: 1) adults manually picked away from progeny, 2) chemical sterilization with FUDR where 100 µL of 10 mg/mL FUDR was dropped onto the food source, and 3) temperature sensitive *glp-4(bn2)* grown at the 22 °C restrictive temperature. Animals were grown on ev from the L1 stage and imaged during day 1, 5, 9, and 13 of adulthood. Imaging was performed for MLS::GFP expressed in the **(A)** muscle, **(B)** intestine, and **(C)** hypodermis.

**Fig. 5.**
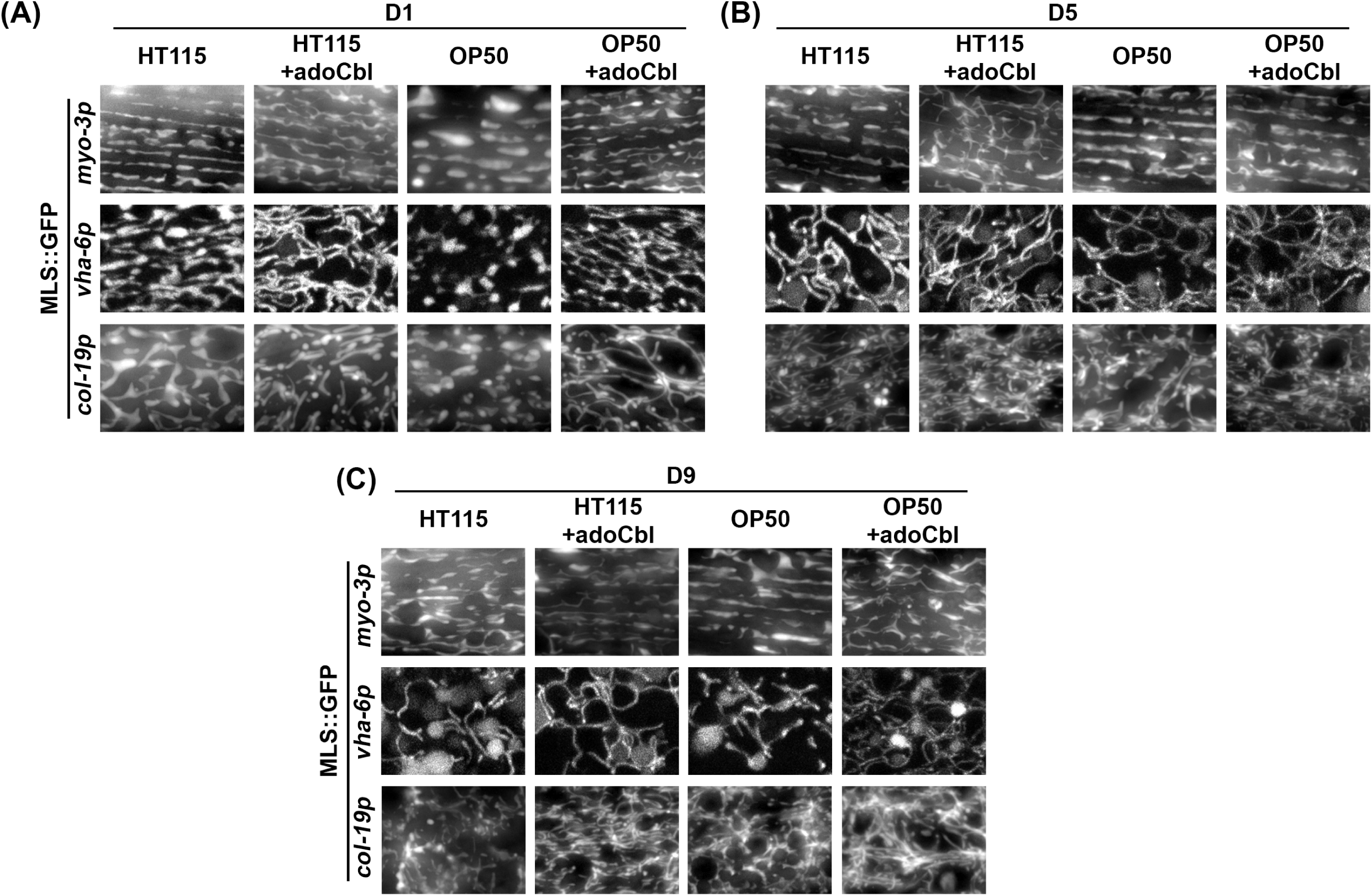
Comparison of mitochondrial morphology on different diets. Animals were grown on an HT115 or OP50 diet with or without the vitamin B12 analog adoCbl. Animals were grown on ev from the L1 stage and imaged during day 1, 5, 9, and 13 of adulthood. Imaging was performed for MLS::GFP expressed in the **(A)** muscle, **(B)** intestine, and **(C)** hypodermis.

### mosSCI generated single-copy MLS::GFP strains exhibit mild physiological changes

Since the most commonly used multi-copy MLS::GFP and TOMM-20::mRFP strains exhibited changes to several physiological measurements including longevity (Valera-Alberni et al, 2024), we next sought to characterize our mosSCI MLS::GFP strains for potential changes in organismal health and longevity. The multi-copy MLS::GFP strains showed minimal changes to lifespan, although one biological replicate out of four total replicates showed a mild decrease in lifespan (**Fig. 6A, Fig. S3**). To further assess animal health, we measured locomotor behavior, and saw no change in thrashing rates throughout aging in any MLS::GFP strains (**Fig. 6B**). Finally, to more carefully evaluate mitochondrial function, we measured oxygen consumption rate (OCR) using a Seahorse assay. Interestingly, we found that all MLS::GFP strains exhibited a lower basal OCR compared to wild-type animals. To ensure that this was due to a reduction in mitochondrial respiration, we measured OCR after treatment with sodium azide, which completely shuts down mitochondrial respiration and saw no difference between the MLS::GFP strains and a wild-type control, suggesting that the decrease in OCR is due to a decline in mitochondrial respiration. Overall, our data show that our MLS::GFP strains are not completely benign and may have a mild impact on mitochondrial respiration, but do not dramatically impact longevity or healthspan unlike the previously developed multi-copy strains (Valera-Alberni et al, 2024).

**Fig. 6.**
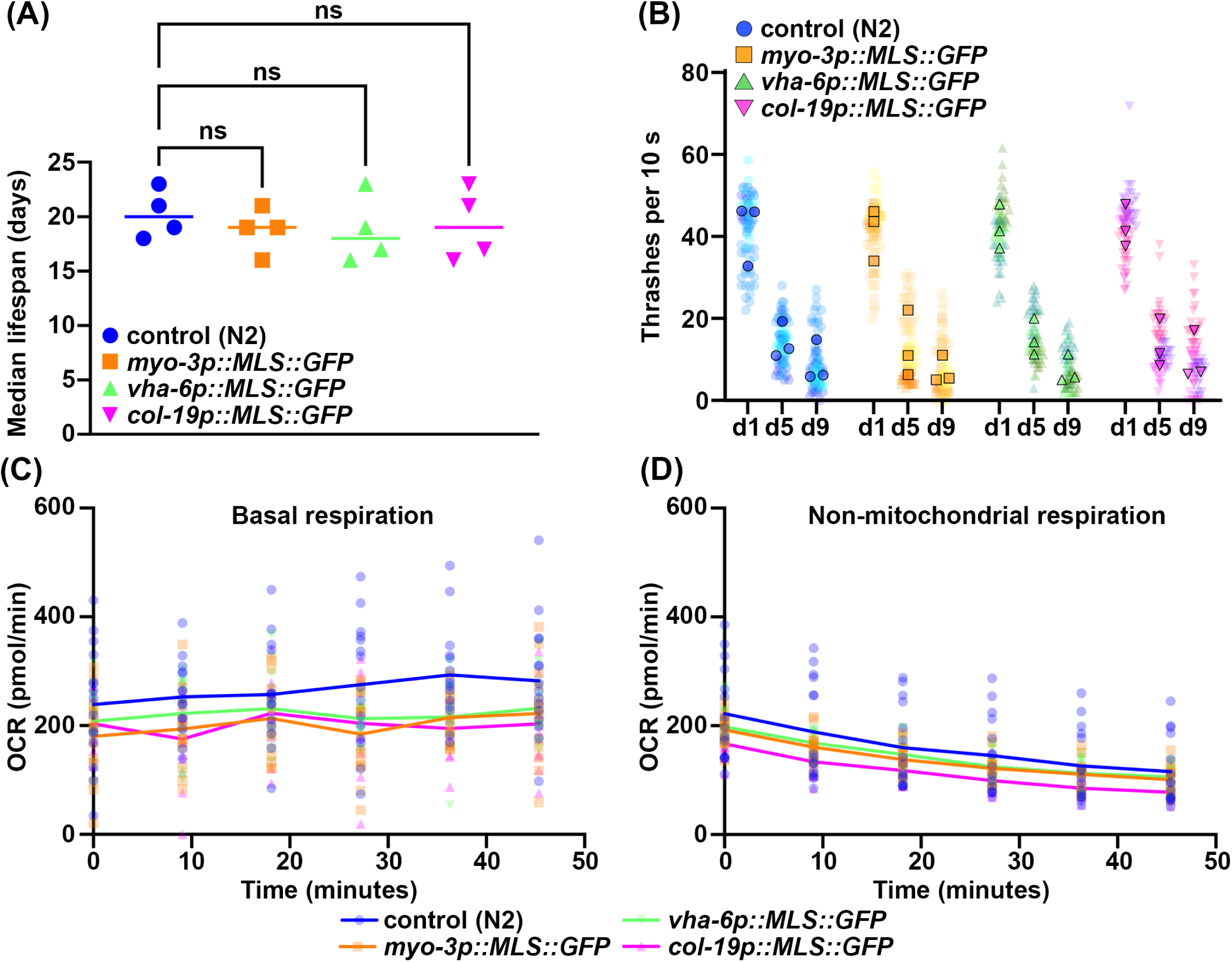
Expression of MLS::GFP has very mild impacts on animal physiology. (**A**) Median lifespans of N2, *myo-3p::MLS::GFP, vha-6::MLS::GFP, col-19p::MLS::GFP*. Each dot represents the median lifespan of a replicate with worms of n > 90. (N=4) (**B**) Thrashing measurements were performed on N2, *myo-3p::MLS::GFP, vha-6::MLS::GFP, col-19p::MLS::GFP* grown on ev from the L1 stage and scored on day 1, 5, and 9 of adulthood. n=20 for each strain per replicate. N=3. All three replicates are superplotted and the mean values of each replicate are indicated with outlined symbols. (**C**) Basal respiration of n > 50 worms measured at day 1 of adulthood across six timepoints in 9 min interval in M9 solution. N=3. (**D**) Non-mitochondrial respiration of n > 50 worms measured at six timepoints after 50mM sodium azide treatment in 9 min interval. N=3. Statistics by one-way ANOVA, using GraphPad Prism 10.0. ns = not significant, * = p < 0.03; ** = p <0.002; *** = p<0.0002; **** = p< 0.0001.

## Discussion

Imaging of mitochondrial morphology is a robust and simple method to get a general idea of mitochondrial quality, as changes to morphology are often correlated with changes to numerous functional measurements of mitochondria (Sharma et al, 2019; Chen et al, 2023). *C. elegans* serve as a robust model to perform mitochondrial imaging during aging, as its short lifespan and small, clear body allows for imaging of mitochondrial morphology throughout the entire lifespan of the worm in adult animals. However, there are many different methods to image mitochondrial morphology in the worm, each with its distinct advantages and disadvantages. Conventional mitochondrial dyes like MitoTracker and TMRE are great options since they do not require strain construction, but suffers from variability in amount of staining across cells, tissues, and individual animals, especially in the *C. elegans* model where the thick cuticle prevents entry of many dyes (Presley et al, 2003; Wang et al, 2016; Valera-Alberni et al, 2024). To circumvent this issue, researchers can deliver these dyes through their bacterial food source, but this method will not allow for robust or equal staining across all tissues (Ravi et al, 2021; Valera-Alberni et al, 2024). These caveats combined with the ease of genetic manipulation in the worm make genetically encoded fluorescent protein-based imaging strategies the most commonly used tools. However, even amongst fluorescent protein-based imaging, there are many different strains, each with individual advantages and disadvantages.

Here, we present single-copy, matrix-localized GFP using a GFP bound to the MLS of ATP-1. Importantly, these transgenes were introduced using mosSCI technology into a known genetic locus, thus preventing unwanted off-target effects of irradiation-based integration methods that integrate into an unknown locus and may interfere with expression of important genes (Thellmann et al, 2003; Frøkjær-Jensen et al, 2008). In addition, we show that our low-copy constructs have limited effects on mitochondrial function and organismal health with only minor effects on OCR, unlike the high-copy expression strains that have numerous effects on mitochondrial function and have been shown independently by another lab to significantly affect whole organism physiology (Valera-Alberni et al, 2024). The MLS::GFP strains are complementary to the membrane-targeted fluorophores published recently by the Mair lab while this manuscript was in preparation. The Mair lab used CRISPR-Cas9 technology to either endogenously tag mitochondrial membrane-localized proteins or express fluorophores with a minimal MLS of outer membrane proteins, and thus have similar benefits to our strains of single-copy expression in known genetic loci. Each strain also presents its own unique advantages, where the Mair lab strains allow for visualization of mitochondria across the entire animal as it is ubiquitously expressed, whereas our strains allow for focusing on a single tissue as we utilized cell-type specific promoters. The Mair lab constructs allow for visualization of membranes, which has much higher resolving capacity to look at mitochondrial substructures, whereas matrix localized fluorophores allow for accumulation of fluorescence in one area and thus is often brighter. Each strain can be utilized for fluorescence recovery after photobleaching (FRAP) experiments where our strains will allow for measurements of mitochondrial matrix continuity, while the Mair lab strains are optimal for measuring membrane fluidity and dynamics. Finally, we also show that these strains can be used together to simultaneously visualize the matrix and outer membrane.

Even amongst the strains presented in both studies, there are considerations to be made. The Mair lab found that red fluorophores – regardless of their identity (i.e., mCherry, mScarlet, mRFP) – showed aggregation in the mitochondria at old age. We confirmed these findings using our matrix-localized mRuby construct, adding further evidence that this aggregation is common across multiple red fluorophores and that it is not limited to just mitochondrial membranes. This is an important consideration as aggregation of proteins both inside and at the outer membrane of mitochondria can result in induction of mitochondrial stress (Berendzen et al, 2016). Thus, some researchers may choose to avoid red fluorophores when regulation of mitochondrial protein homeostasis is the primary area of study. In addition to red fluorophores, blue fluorescent proteins have commonly been shown to have problems for mitochondrial imaging and perturb cellular health (Higuchi-Sanabria et al, 2016), potentially due to high levels of ROS production (Alvarez et al, 2010). While some labs have prioritized identification of non-toxic blue fluorophores (Mohapatra et al, 2019), they have yet to be tested in the *C. elegans* system.

Although we focused primarily on strain choice in this manuscript, there are also many additional important considerations for imaging of mitochondrial morphology. First, bacterial food choice is critical as previous reports have shown that mitochondrial morphology may be different when animals are grown on the two most standard bacterial food choices, OP50 and HT115, due to deficiencies in vitamin B12 (Revtovich et al, 2019). The K strain *E. coli* HT115 is most commonly used for RNAi as the two largest RNAi libraries – the Vidal and Ahringer libraries – were constructed in this RNAi competent strain (Ahringer, 2006). Subsequently, modifications were made to the B strain *E. coli* OP50 to make them RNAi competent by deleting the RNAIII RNase and genomically introducing an IPTG-inducible T7 RNA polymerase (Neve et al, 2020). However, no thorough RNAi library exists yet in this construct and utility of this strain requires cloning each individual gene of interest into an RNAi vector and transforming it into this modified OP50 strain. Thus, for large-scale RNAi studies, the usage of HT115 is still unavoidable. However, our studies show that during aging, there are not major differences between the OP50 and HT115 diet in terms of mitochondrial morphology; simply the OP50 diet displays slightly more fragmentation of mitochondria, but the trends for age-associated fragmentation are robustly apparent in either diet. Moreover, as previously shown, mitochondrial morphology can be matched between OP50 and HT115 diets, solely by addition of vitamin B12 in the OP50 diet, and we’ve found this to be true across the lifespan of the animal. Thus, unless major metabolic pathways are being tested where either excess vitamin B12 or other still uncharacterized differences between OP50 and HT115 diets may cause unmanageable confounding variables, we believe that standard mitochondrial imaging during aging is not extremely sensitive to differences in these standard diet choices. However, care should be taken by the researcher to confirm this for each of their experimental conditions.

Beyond metabolic differences between OP50 and HT115, researchers should also consider the usage of antibiotics. For HT115 bacteria, growth on tetracycline is often used to select for the correct bacteria as the RNase III allele (*rnc:14*::Δ*Tn10*) in HT115 bacteria confers tetracycline resistance (Neve et al, 2020). Moreover, the pL4440 vector often used as the expression vector for dsRNA for RNAi experiments carries an ampicillin resistance gene, which researchers often select for using either ampicillin or the more shelf-stable carbenicillin. However, previous research has shown that exposure to specific antibiotics – including tetracycline (Chatzispyrou et al, 2015) and ampicillin (Khan et al, 2014) – can impact mitochondrial function. While these studies may argue that usage of antibiotics should be avoided, this can be challenging in some cases as RNAi requires selection of plasmid-carrying bacteria and some common lab contaminants have been shown to impact many criteria of organismal health and longevity (Stuhr & Curran, 2023). Importantly, our study has already highlighted how OP50 and HT115 conditions are not dramatically different, especially when vitamin B12 differences are corrected for. Since our HT115 growth conditions include exposure of worms to both carbenicillin and tetracycline while OP50 growth conditions do not, we can likely extrapolate these data to suggest that there are not major concerns for using antibiotics for the specific mitochondrial imaging conditions used in this study.

To further add to complications, other technical components can impact mitochondrial imaging. For example, a standardized method needs to be used to synchronize and age *C. elegans* populations. The most common method of synchronization is to perform an egg prep by bleaching of animals using a sodium hypochlorite solution; however, this applies significant stress to the animals, which can impact metabolism (Verdú et al, 2022). As an alternative, egg-lay methods (Castro Torres et al, 2022) or manually picking animals at definable stages such as visibility of the L4 “crescent”-shared pre-vulva, but these assays are generally manually intensive and would be challenging for large-scale assays. As an alternative, a commercially available *C. elegans* synchronizer can be used to harvest large volumes of L1 animals from a mixed population (Rasmussen & Reiner, 2021), but this may be cost-prohibitive for some labs. To compound on this issue, once a synchronized population is established, care must be taken to identify a method of choice for aging out cohorts of animals. Here, we provide three methods to age out animals: first, animals can be manually manipulated away from progeny. While this is the most “natural” method to age out animals and do not require any interventions, this is also the most manually intensive and is more challenging for large-scale experiments. We also tested both chemical and genetic sterilization techniques and found that they do not dramatically change mitochondrial morphology with age, although the timing may shift slightly. This shift in timing can be due to technical aspects, such as lack of manual manipulation of worms when using FUDR, which can reduce physical stress to the animals and delay mitochondrial fragmentation. Or in the case of using a temperature-sensitive mutant, the elevated temperatures may serve as a stress on the worm that can accelerate mitochondrial aging.

Overall, there are many considerations to be made when performing mitochondrial imaging in a laboratory, particularly during the aging process. Therefore, care must be taken to standardize methods for mitochondrial imaging in each laboratory, or proper controls must be performed using multiple methods to confirm that phenotypic findings are not artifacts of methods. Finally, while mitochondrial morphology can be used as an indirect measurement of mitochondrial function since morphology often correlates with mitochondrial function, there are many exceptions to this correlation (Osellame et al, 2012). Therefore, to perform a comprehensive analysis of mitochondrial health and function, additional measurements need to be made, including measurements of mitochondrial membrane potential (Chen, 1988; Sakamuru et al, 2016), ATP synthesis capacity (Zong et al, 2024), calcium levels (Pivovarova & Andrews, 2010), respiratory capacity (Zong et al, 2024), and mitochondrial DNA sequence and content (Castellani et al, 2020). However, as these assays can be technically challenging, imaging of mitochondrial morphology can be used as a first step in determining whether any experimental conditions affect general mitochondrial biology.

---

**Fig. S1.**
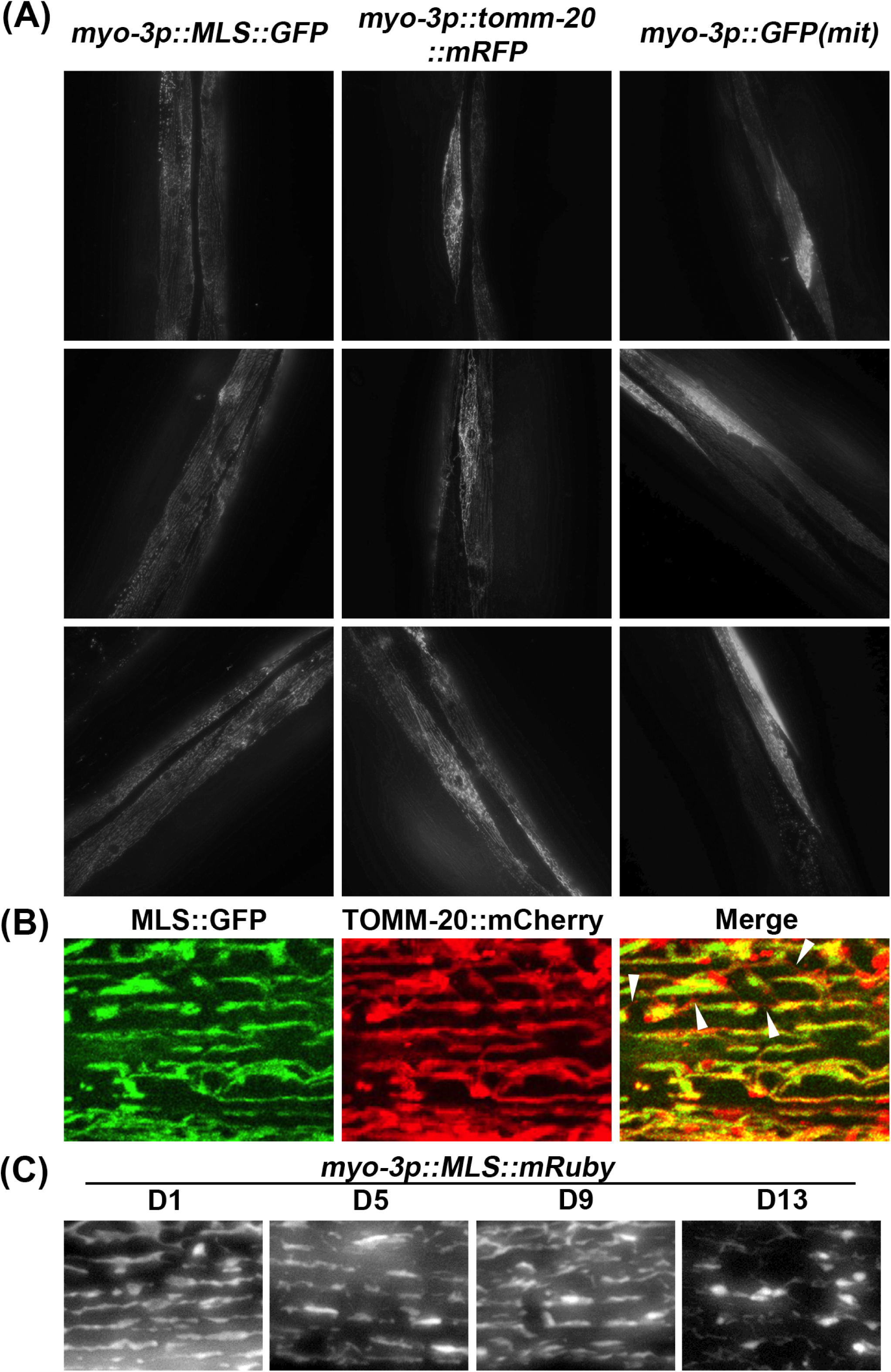
Comparative imaging of muscle mitochondria across strains. **(A)** Comparison of day 1 imaging of mosSCI MLS::GFP (RHS191) strain to multi-copy *myo-3::GFP(mit)* (SJ4103) and multi-copy *myo-3p::TOM20::mRFP* (PS6192) showing variability in SJ4103 and PS6192 but not in RHS191. Images are intentionally shown zoomed out to display multiple muscle cells per image. (**B**) Imaging of muscle MLS::GFP x TOMM-20::mCherry from the Mair lab at day 5 of adulthood. Arrowheads indicate outer membrane mCherry signal in the absence of matrix MLS::GFP. (**C**) Imaging of muscle MLS::mRuby during aging. All animals were grown on ev from the L1 stage and imaged at the indicated days.

**Fig. S2.**
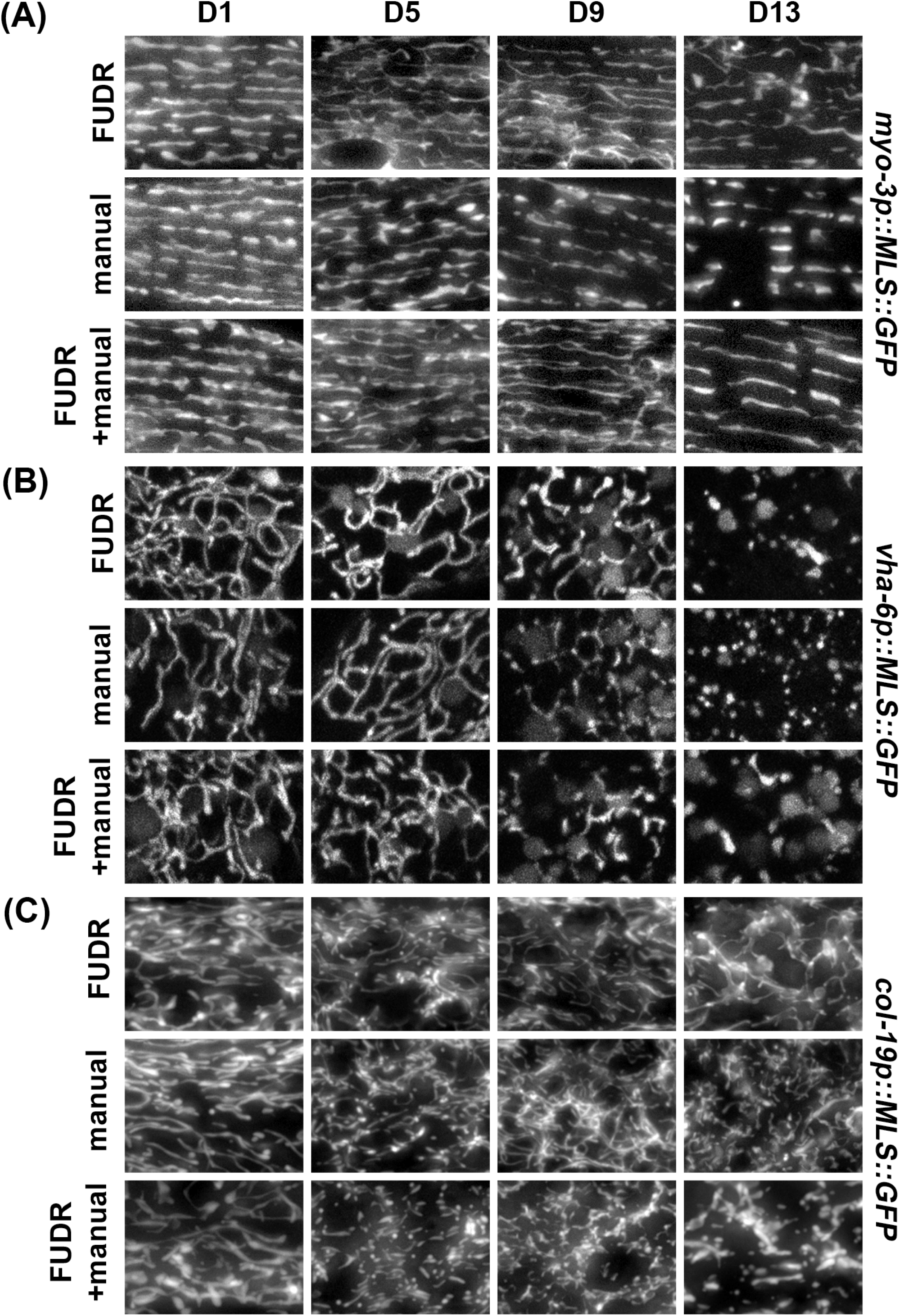
Comparison of manual manipulation of worms versus no manipulation of worms. Animals expressing tissue-specific MLS::GFP were grown on ev from the L1 stage. At day 1 of adulthood, animals were either moved onto plates supplemented with 100 µL of 10 mg/mL FUDR directly onto the food source, or on standard RNAi plates. Animals on standard RNAi plates and FUDR plates were moved daily (manual and FUDR+manual) or left undisturbed (FUDR). Imaging was performed at day 1, 5, 9, and 13 of adulthood in the **(A)** muscle, **(B)** intestine, and **(C)** hypodermis.

**Fig. S3.**
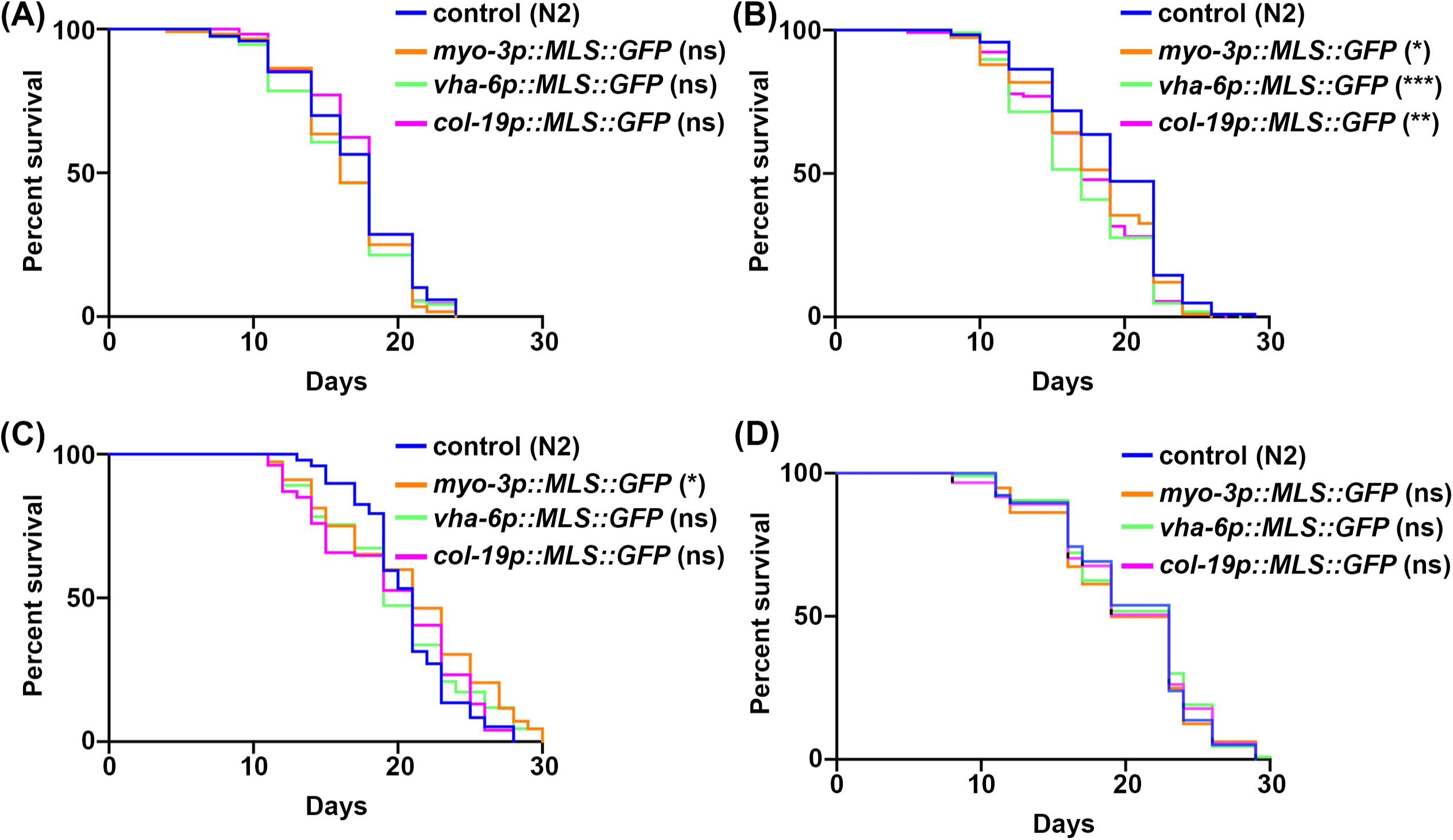
MLS::GFP expression only has mild impacts on lifespan. Lifespans were performed on wild-type control (N2), and strains expressing MLS::GFP in the muscle (*myo-3p*), intestine (*vha-6p*), and hypodermis (*col-19p*), grown on ev from the L1 stage. All replicates with worms n > 90. Graphs were plotted and statistically analyzed by log-rank test (Mantel-Cox), using GraphPad Prism 10.0. ns = not significant, * = p < 0.03; ** = p <0.002; *** = p<0.0002; **** = p< 0.0001. All lifespan statistics are available in **Table S2**.

**Table S1. Table of lifespan statistics.**

## Materials and Methods

### C. elegans strains and maintenance

All strains used in this study are derived from the N2 wild-type animal from the Caenorhabditis Genetics Center (CGC) and are listed in **Table 1**. Animals are maintained at 15 °C on OP50 *E. coli* B strain bacteria on standard NGM plates (Nematode Growth Medium, 1 mM CaCl_2_, 5 µg/mL cholesterol, 25 mM KPO_4_, 1 mM MgSO_4_, 2% agar w/v, 0.25% Bacto-Peptone w/v, 51.3 mM NaCl) plates. Animals are maintained by either chunking a small patch of worms or manually picking a small population of young (before L4) animals onto freshly seeded plate. Animals are only kept for approximately 25-30 generations in this way before thawing a new batch to avoid genetic drift.

**Table 1.**

| Strains used in this study |  |  |
| --- | --- | --- |
| *C. elegans*: Bristol (N2) strain as wild type (WT) | CGC | N2 |
| *C. elegans*: RHS19: *glp-4(bn2)* | CGC | SS104<br>backcrossed 6x |
| *C. elegans*: RHS191: *uthSi17[myo-3p::MLS::GFP::unc-54 3'UTR::cb-unc-119(+)] I; unc-119(ed3) III* | (Moehle et al, 2021) | AGD1664<br>backcrossed 4x |
| *C. elegans*: RHS192: *uthSi83[col-19p::MLS::GFP::unc-54 3'UTR::cb-unc-119(+)] I; unc-119(ed3) III* | (Moehle et al, 2021) | AGD2837<br>backcrossed 4x |
| *C. elegans*: RHS193: *uthSi80[vha-6p::MLS::GFP::unc-54 3'UTR::cb-unc-119(+)] I; unc-119(ed3) III* | (Moehle et al, 2021) | AGD2805<br>backcrossed 4x |
| *C. elegans*: RHS213: *zcls[ges-1p::GFP(mit)]* | CGC | SJ4313<br>backcrossed 3x |
| *C. elegans*: RHS218: *uthSi17[myo-3p::MLS::GFP::unc-54 3'UTR::cb-unc-119(+)] I; unc-119(ed3) III; glp-4(bn2)* | This study |  |
| *C. elegans*: RHS243: *uthSi83[col-19p::MLS::GFP::unc-54 3'UTR::cb-unc-119(+)] I; unc-119(ed3) III; glp-4(bn2)* | This study |  |
| *C. elegans*: RHS217: *uthSi80[vha-6p::MLS::GFP::unc-54 3'UTR::cb-unc-119(+)] I; unc-119(ed3) III; glp-4(bn2)* | This study |  |
| *C. elegans*: RHS218: *glp-4(bn2) I; uthSi17[myo-3p::MLS::GFP(65C)::unc-54 3'UTR, cb-unc-119(+)] I;* | This study |  |
| *C. elegans*: RHS180: *wbmls98[eft-3p::tomm-20(aa1-49)::mCherry::unc-54 3'UTR]; wbmls65[eft-3p::3XFLAG::dpy-10 crRNA::unc-54 3'UTR]; uthSi17[myo-3p::MLS::GFP(65C)::unc-54 3'UTR, cb-unc-119(+)] I;* | This study |  |
| *C. elegans*: PS6192: *syIs243[myo-3p::TOMM-20::mRFP + unc-119(+)] + pBS Sk+* | CGC |  |
| *C. elegans*: SJ4103: *zcls14[myo-3p::GFP(mit)]* | CGC |  |
| *C. elegans*: SJ4143: *zcls17[ges-1p::GFP(mit)]* | CGC |  |
| *C. elegans*: AGD2319: *unc-119(ed3) III*;<br>*uthSi62[vha-6p::MLS::mRuby::unc-54 3'UTR, cb-unc-119(+)] IV*; | (Tharp et al, 2021) |  |
| *C. elegans*: AGD2883: *unc-119(ed3) III*;<br>*uthSi90[myo-3p::MLS::mRuby::unc-54 3'UTR cb-unc-119(+)] IV*; | This study |  |
| *C. elegans*: RHS243: *uthSi83[col-19p::MLS::GFP(65C)::unc-54 3'UTR, cb-unc-119(+)] I*; *glp-4(bn2) I*; | This study |  |

For all experimental purposes, animals are age-matched using a standard bleaching protocol as previously described (Bar-Ziv et al, 2020). Briefly, animals are collected using M9 solution (22 mM KH_2_PO_4_ monobasic, 42.3 mM Na_2_HPO_4_, 85.6 mM NaCl, 1 mM MgSO_4_) and bleached using a 1.8% sodium hypochlorite and 0.375 M KOH solution. After bleaching animals, eggs are washed 3-4x with M9 solution with repeated centrifugation at 1,100 x g and aspiration of solution. Intact eggs were floated in M9 solution in a rotator overnight at 20 °C to obtain tighter synchronization at the L1 stage. Synchronized L1 animals were subsequently plated on RNAi plates (1 mM CaCl_2_, 5 µg/mL cholesterol, 25 mM KPO_4_, 1 mM MgSO_4_, 2% agar w/v, 0.25% Bacto-Peptone w/v, 51.3 mM NaCl, 1 µM IPTG, and 100 µg/mL carbenicillin; HT115 *E. coli* K strain containing pL4440 vector control or pL4440 with RNAi of interest) unless otherwise noted. All aging experiments were performed on plates supplemented with 100 µL of 10 mg/mL FUDR spotted directly on the bacterial lawn unless otherwise noted.

For growth of *glp-4(bn2)* animals, we grow animals at 22 °C from the L1 stage. Although previous reports have shown that *glp-4(bn2)* animals are sterile at 25 °C (Beanan & Strome, 1992), after backcrossing animals 6x to our N2 animals we found that our *glp-4(bn2)* animals were fully sterile at 22 °C. Therefore, we opted to grow animals at 22 °C to reduce caveats of potential induction of stress at 25 °C (Gouvêa et al, 2015).

For vitamin B12 supplementation assays, NGM plates were supplemented with 12.8 nM of adenosylcobalamin, vitamin B12 analog. Adenosylcobalamin was added into the media post-autoclaving. Animals were grown on adenosylcobalamin containing plates from the L1 stage throughout their lifespan.

### Making C elegans transgenic strains

Transgenic *C. elegans* strains were generated using the Mos1-mediated Single Copy Insertion (MosSCI) technique, following the detailed protocol described by Garcia et al (Garcia et al, 2022). Specifically, the transgenic strains used in this study were created by injecting MosSCI-specific strains with a plasmid cocktail. This cocktail included plasmid vector with transgene for Mos1 transposes (pCFJ601), transgenic construct designed for tissue-specific expression of MLS::GFP or MLS::mRuby within a MosSCI-compatible vector, and multiple tissue-specific co-injection fluorescent markers. Details of the injection cocktail and MosSCI-specific strains are provided in **Table 2** and **Table 3** respectively.

**Table 2:** MosSCI injection cocktail.

| Plasmid | Transgene | Function | Expression tissue | Working Concentration |
| --- | --- | --- | --- | --- |
| pCFJ601 | eft-3p::Mos1 transposase | Transposase | Ubiquitous | 50 ng/uL |
| pGH8 | rab-3p::mCherry::unc-54 UTR | Co-injection marker | Pan-neuronal | 10 ng/uL |
| pCFJ90 | myo-2p::mCherry::unc-54 UTR | Co-injection marker | Pharynx | 2.5 ng/uL |
| pCFJ104 | myo-3p::mCherry::unc-54 UTR | Co-injection marker | Body wall muscle | 5 ng/uL |
| pCFJ35X | construct, Cb-unc-119(+) | MosSCI construct | Construct dependent | 25 ng/uL |

**Table 3:** mosSCI strains used in this study.

| Strain name | Genotype | promoter | mossci vector | mls gene | MosSCI strain | Integration site |
| --- | --- | --- | --- | --- | --- | --- |
| RHS191 | uthSi17[myo-3p::MLS::GFP::unc-54 3'UTR, cb-unc-119(+)] I; unc-119(ed3) III; | myo-3p | pCFJ352 | atp-1 | eg6701 | Chromosome I |
| AGD2883 | unc-119(ed3) III; uthSi90[myo-3p::MLS::mRuby::unc-54 3'UTR cb-unc-119(+)] IV; | myo-3p | pCFJ356 | atp-1 | eg6703 | Chromosome IV |
| RHS193 | uthSi84[vha-6p::MLS::GFP(65C)::unc-54 3'UTR, cb-unc-119(+)] I; unc-119(ed3) III; | vha-6p | pCFJ352 | atp-1 | eg6701 | Chromosome I |
| RHS192 | uthSi83[col-19p::MLS::GFP(65C)::unc-54 3'UTR, cb-unc-119(+)] I; unc-119(ed3) III; | col-19p | pCFJ352 | atp-1 | eg6701 | Chromosome I |

Additionally, the transgenic MLS::GFP *C. elegans* strains were crossed with *glp-4(bn2)* mutant animals. The *glp-4(bn2)* mutation in the resulting strains were confirmed through sequencing. Single-worm lysis was performed by placing single worms into dH_2_0, proteinase K, and PCR buffer (we used Q5 PCR buffer) at a 42:3:5 ratio. The worm mixtures were then heated to 60 °C for 1 h and 98 °C for 20 min. 1 µL of this lysate was used as template DNA for a standard PCR using forward primer tgacataccattgaggcttgag and reverse primer gtaaattgaccttggttgaggc. Standard sanger sequencing was performed at Genewiz using the forward primer.

### C. elegans microscopy

Imaging of mitochondrial morphology was performed by using either a Leica Thunder microscope equipped with a 63x/1.4 Plan AproChromat objective, standard GFP and dsRed filter, Leica DFC9000 GT camera, a Leica LED5 light source, and run on LAS X software, or Leica Stellaris confocal microscope equipped with a white light laser source and spectral filters, HyD detectors, 63x/1.4 Plan ApoChromat objective, and run on LAS X software. Animals were placed in M9 solution directly on a glass slide, a cover slip is applied, and imaging is performed within 10 minutes of slide preparation. Quantification of mitochondrial morphology is performed using mitoMAPR (Zhang et al, 2019).

### C. elegans lifespan

All lifespan assays were performed on standard RNAi plates with HT115 bacteria at 20 °C as previously described (Castro Torres et al, 2022). Animals were exposed to FUDR from the day 1 adult stage to eliminate progeny. Viability were scored every other day until all animals are scored or censored. Censorship is defined as animals that exhibit deaths unrelated to aging: vivipary (bagging), desiccation on the walls of the petri dish, intestinal leakage out of the vulva, etc. Survival curves were plotted and LogRank statistical analyses were performed using Prism software. All statistical data for lifespans are available in **Table S1**.

### C. elegans seahorse assay

Seahorse assay was performed in day 1 adult animals synchronized using a standard bleaching protocol. Animals were collected off plates using M9, bacteria were washed with M9 solution 3x using repeated centrifugation/aspiration. ∼10-15 worms were pipetted into each well of a Seahorse XF96 cell culture microplate. Basal oxygen consumption rate was measured using an Fe96 sensor cartridge on a Seahorse XFE96 Analyzer with 3 minutes mixing, 2 minutes wait, and 2 minutes measuring. Non-mitochondrial respiration rates were measured using 50mM sodium azide as previously described (Haroon & Vermulst, 2019; Ng & Gruber, 2019). Oxygen consumption rate was normalized for number of worms.

### Statistical analyses

For all imaging experiments, quantification was performed using mitoMAPR and statistical analysis was performed using one-way ANOVA statistical testing. For lifespans, LogRank testing (Mantel-Cox) was performed. For seahorse analysis, one-way ANOVA testing was used. All statistical tests were performed using Prism software. All experiments were performed across a minimum of 3 biological replicates.

## Acknowledgements

J.K. is supported by the USC Graduate Provost Fellowship; M.V. is supported by 1R25AG076400; A.B. is supported by the USC Undergraduate Provost Fellowship; M.A. and G.G. are supported by T32AG052374; and R.H.S. is supported by R01AG079806 from the National Institute on Aging and 2022-A-010-SUP from the Larry L. Hillblom Foundation. Some strains were provided by the CGC, which is funded by the NIH Office of Research Infrastructure Programs grant P40 OD010440. Some gene analysis was performed using Wormbase, which is funded on a U41 grant HG002223.

